# Transcriptome Profiling and Characterization of Peritoneal Metastasis Ovarian Cancer Xenografts in Humanized Mice

**DOI:** 10.1101/2023.10.27.563867

**Authors:** Sung Wan Kang, Ji-young Lee, Ok-Ju Kang, Yong-Man Kim, Eun Kyung Choi, Shin-Wha Lee

## Abstract

**Background:** Although immunotherapy has not yet been as successful in ovarian cancer (OC), it remains a potential therapeutic strategy. Preclinical models of OC are necessary to evaluate the efficacy of immuno-oncology (IO) drugs targeting human cancer and immune components but have been underutilized. Developing mouse models with a humanized (Hu) immune system can help understand the human immune response to IO drugs, including immune checkpoint inhibitors (ICIs), which have demonstrated limited effectiveness in OC patients.

**Methods:** We established OC xenograft Hu-mouse models by intraperitoneally injecting luciferase-expressing SKOV-3 Luc and OVCAR-3 Luc OC cells into CD34^+^ Hu-mice. Tumor growth was monitored through bioluminescence imaging (BLI). We assessed the efficacy of PD-1 blockade with pembrolizumab in the SKOV-3 Luc Hu-mouse model. The immune profiles of the tumors were characterized using colorimetric immunostaining and flow cytometry. Additionally, we analyzed RNA-seq data to investigate the gene expression signature of pembrolizumab refractory tumors.

**Results:** We confirmed tumor development in both OC cell lines within CD34^+^ Hu-mice. In these models, human lymphocyte and myeloid cell subsets were present in the tumors, draining lymph nodes, blood, and spleens. The SKOV-3 Luc tumor-bearing Hu-mice did not respond to pembrolizumab monotherapy. These tumors exhibited a high presence of tumor-infiltrating macrophages. Tumors in Hu-mice unresponsive to pembrolizumab showed a lower abundance of CD8^+^ T-cells, memory B cells, plasma cells, and a higher proportion of naïve M0 macrophages and mast cells compared to the PBS control. Furthermore, we identified 43 significantly enriched gene sets in these tumors. The differentially expressed genes (DEGs) were predominantly enriched in HDAC class I, RB1, KLF1/3, TCF21, MYD88, SMARCE1 target genes, and genes associated with epithelial-mesenchymal transition (EMT) and fibroblasts.

**Conclusion:** Our xenograft Hu-mouse model of OC provides a valuable tool for investigating the efficacy of IO drugs. The insights gained from this model offer potential avenues to explore mechanisms of resistance to PD-1/PD-L1 blockade in OC.

## INTRODUCTION

Ovarian cancer (OC) is the deadliest gynecologic malignancy. Approximately 70% of OC cases are diagnosed at an advanced stage due to the lack of distinct symptoms before metastasis to distant regions [1, 2]. The standard treatment for OC typically comprises debulking surgery and chemotherapy involving platinum and taxane, with the recent addition of poly (ADP-ribose) polymerase (PARP) inhibitors. Despite an initial positive response to these standard treatments in patients with advanced-stage OC, recurrence is frequent, resulting in a 5-year survival rate of less than 50% [3, 4].

Immunotherapy has emerged as a promising therapeutic intervention for cancer treatment. Approaches such as immune checkpoint inhibitors (ICIs) therapy, infusion of tumor-infiltrating lymphocytes (TILs), mRNA vaccination, and adoptive T-cell therapies using chimeric antigen receptors (CARs) or engineered T-cell receptors (TCRs) are being explored in various cancer patients [5, 6]. Notably, the PD-1/PD-L1 blockade has significantly improved overall survival in certain cancers such as non-small cell lung cancer and melanoma [7, 8]. OC is generally classified as an immunogenic tumor, and an increase in CD8^+^ T-cell abundance has been associated with improved survival rates among OC patients [9]. However, despite the immunogenicity of OC, clinical trial results suggest that OC patients may not benefit from ICI therapy [10]. This ineffectiveness could be attributed to the highly immunosuppressive nature of the tumor microenvironment (TME), low mutational burden, and limited T-cell infiltration in these patients [11, 12]. Therefore, there is a pressing need for additional therapeutic strategies to effectively treat these individuals. To address the issue of low response rates, it is crucial to understand the complexities of the immunological response. This understanding will facilitate the identification of novel combination immunotherapy regimens and predictive biomarkers for treatment effectiveness, thereby enabling personalized immunotherapy through patient stratification.

The development of new immunotherapy strategies for OC, given its tumor heterogeneity, is hindered by the lack of readily available, high-precision preclinical tumor models. The use of in vivo preclinical models is essential for determining the efficacy of potential drug candidates before their introduction into clinical trials. Syngeneic models, such as the widely used ID8 orthotopic murine model and its genetically modified variants, have been extensively used to study the role of the immune system in ovarian cancer progression and to analyze therapeutic responses [13-15]. However, these syngeneic models do not accurately represent human disease. This discrepancy becomes problematic when the therapeutic agent under investigation is responsive only to human isoforms, or when the expression levels of the targets of interest vary between human and mouse immune cell subsets. Therefore, there is a need for comprehensively characterized mouse models with a functionally intact humanized immune system. These models are crucial for assessing immunotherapy agents and understanding the underlying mechanisms contributing to the lack of response to immunotherapy.

Humanized (Hu) mouse models, featuring cell line-derived xenograft (CDX) or patient-derived xenograft (PDX) tumors, have been utilized to study the efficacy of PD-1/PD-L1 immune checkpoint inhibitors (ICIs) across various cancer types. These models are established in immunodeficient mice engrafted with CD34^+^ hematopoietic stem cells (HSCs), allowing for the characterization of distinct circulating and tumor-infiltrating immune cell profiles, as well as the observation of diverse anti-tumor efficacy of PD-1/PD-L1 ICIs [16-18]. Specifically for OC research, intraperitoneal and subcutaneous xenograft models of defined OC cell lines and patient-derived cells have been employed. These models are used to study the effect of immunotherapy agents in Hu-mice, as well as cytotoxic therapeutics and targeted agents in immunocompromised mice, such as nude and NOD/SCID/IL2rγnull (NSG) mice [19-22].

In this study, we established and characterized two intraperitoneal CDX models of OC in Hu-mice engrafted with CD34^+^ HSCs, and evaluated their responsiveness to PD-1 blockade. Our investigation centered on changes in the TME during anti-PD-1 treatment for OC. To this end, we examined the cellular and molecular characteristics of xenografts in CD34^+^ Hu-mice undergoing PD-1 blockade treatment.

## MATERIALS AND METHODS

### Cell lines

The SKOV-3-Luc-D3 cell line was purchased from Caliper Life Sciences (MA, USA). The OVCAR-3 cell line was obtained from the Korean Cell Line Bank (KCLB, Seoul, Korea) and transfected with the pGL3 vector (WI, USA) containing the firefly luciferase gene. Cell lines were grown in RPMI-1640 medium supplemented with 10% fetal bovine serum and 1% antibiotic-antimycotic solution at 37°C.

### Mice

For this study, 20-weeks-old female NOD.Cg-Prkdc^scid^Il2rγ^tm1Wjl^/SzJ (NSG) and NOD/ShiLtJ-Prkdc^em1AMC^ Il2rg^em1AMC^ (NSGA) mice engrafted with human CD34^+^ HSCs were obtained from Jackson Laboratory (ME, USA) and JA BIO (Suwon, Korea). When the mice were 3-4 weeks old, they were exposed to whole-body irradiation. Subsequently, each mouse received an intravenous injection of 1 × 10^5^ CD34^+^ HSCs. All animal experiments were approved by the Institutional Animal Care and Use Committee (IACUC) of the Asan Institute for Life Sciences at the Asan Medical Center (approval ID: 2022-12-233), and the studies were conducted in accordance with the approved guidelines and regulations. All mice were bred and maintained under a 12:12 light–dark cycle in the experimental facility at the disease animal resource center (Seoul, Korea) where received sterilized food and water *ad libitum*.

### In vivo experiments

In the intraperitoneal xenograft model, 1×10^7^ SKOV-3 Luc and OVCAR-3 Luc cells, mixed with 50% matrigel, were injected into the peritoneal cavity of humanized NSG or NSGA mice 17-18 weeks post-transplantation of CD34^+^ cells. Tumor development was tracked on a weekly basis using bioluminescence imaging. For this, mice were anesthetized with isoflurane, followed by an intraperitoneal injection of D-luciferin solution (150 mg/kg). Imaging was conducted using an IVIS Spectrum system 10 minutes post-injection. The bioluminescent signal was quantified using Living Image 3.1 (Caliper Life Sciences, CA, USA). To investigate the sensitivity to PD-1 blockade, mice were imaged using the IVIS system two weeks post-inoculation of SKOV-3 Luc cells to confirm tumor development. They were then randomly divided into two groups of eight: a PBS control group, and a group treated with Pembrolizumab. PBS and pembrolizumab (10 mg/kg) were administered intraperitoneally, twice a week for three weeks. Tumor growth was monitored using the IVIS imaging system. A week after the final treatment, the mice were euthanized, and their tumor tissues were collected for further analysis. Tumor volumes were determined based solely on the tissue where the tumor was identified through histological analysis. Statistical analyses were conducted on these tumor volumes.

### Histological analysis, immunohistochemistry, and immunofluorescence microscopy

After the final in vivo optical imaging, all tumor-bearing mice were humanely euthanized in accordance with our experimental protocol. The tumor tissues located in the peritoneal cavity were isolated, fixed, and embedded in paraffin for subsequent analysis, and the corresponding tissue sections were stained with hematoxylin and eosin (H&E). For the processes of immunohistochemistry (IHC) and immunofluorescence (IF) staining, antigen retrieval was performed by heating the paraffin-embedded tissue sections in Tris-EDTA buffer (Vector Laboratories, CA, USA). Subsequently, the slides were blocked using 2.5% normal goat serum blocking solution (Vector Laboratories, CA, USA) for 30 minutes at room temperature (RT). Endogenous peroxidase activity, relevant for IHC, was inhibited using BLOXALL Endogenous blocking solution (Vector Laboratories, CA, USA). The tissue sections were then stained overnight at 4°C with primary antibodies against anti-CD3 (ab52959, Abcam, CA, USA), anti-CD4 (AF-379-NA, R&D Systems, MN, USA), anti-CD8 (NB100-65729, Novus Biologicals, CO, USA), and anti-PD-L1 (E1L3N, Cell Signaling). These sections were subsequently incubated for 2 hours at RT with secondary antibodies. For IHC, biotinylated mouse or rabbit IgG (Abcam, CA, USA) were used, while for IF, mouse IgG tagged with Alexa Flour 488, goat IgG tagged with Alexa Flour 555, and rabbit IgG labeled with Alexa Flour 647 (Invitrogen, CA, USA) were applied. Further, IHC slides were subjected to a 30-minute incubation at RT with ABC reagents (Vector Laboratories, CA, USA), followed by 10-minute staining with DAB, and counterstaining with 1% methyl green. Finally, the sections were embedded with mounting medium, applicable for IHC and IF (with DAPI), respectively. The images of H&E and IHC were acquired using the BX53 microscopy (Olympus, Tokyo, Japan) and VS200 digital slide scanner (Olympus, Tokyo, Japan). In addition, the IF images were captured using an LSM 710 confocal microscope (Carl Zeiss, Oberkochen, Germany).

### Flow cytometric analysis

Tumor tissues were dissociated for 1 hour at 37°C using RPMI with 10% FBS, 1 mg/ml type IV collagenase (Sigma-Aldrich, MO, USA), 0.2 mg/ml hyaluronidase (Sigma-Aldrich, MO, USA), and 0.1 mg/ml DNase I (Roche, Basel, Switzerland). Single cells were mechanically isolated from lymph nodes and spleen tissues using cell strainers and syringe plungers. Peripheral blood mononuclear cells (PBMCs) were separated using the Histopaque-1077 (Sigma-Aldrich, MO, USA) gradient method. The cell surface was blocked with an Fc-block antibody (BD Biosciences, CA, USA) and subsequently stained with respective fluorochrome-conjugated antibodies for 30 minutes. The antibodies used for analysis were from Biolegend (CA, USA) or Invitrogen (CA, USA), including anti-CD45 (304029, HI30 clone), anti-CD3 (300406, UCHT1 clone), anti-CD4 (300514, RPA-T4 clone), anti-CD8 (344710, SK1 clone), anti-CD56 (318318, HCD56 clone), anti-CD11b (301306, ICRF44 clone), anti-CD19 (302218, HIB19 clone), and anti-PD-1 (12-9969-42, MIH clone). PE Mouse IgG1 kappa isotype control from BD Biosciences (CA, USA) was used in parallel to PD-1. Dead cells were excluded with a Live/Dead Cell Stain kit (Invitrogen, Carlsbad, CA). Single cells were identified by FACS Canto II (BD Biosciences, CA, USA), and the data were analyzed with FlowJo software (TreeStar, OR, USA).

### RNA-seq analysis

Total RNA was isolated from the tumor tissues of SKOV-3 Luc xenograft in Hu-mice treated with either PBS or pembrolizumab. The isolation was performed using the mirVana miRNA isolation kit (Thermo Fisher Scientific, MA, USA). A library was prepared with 100 ng of total RNA for each sample using the TruSeq RNA Access library prep kit (Illumina, CA, USA). Subsequently, the indexed libraries were submitted to an Illumina NovaSeq (Illumina, CA, USA), and the paired-end (2×100 bp) sequencing was performed by Macrogen.

### Data processing and differential gene expression analysis

Trimmomatic v0.38 was utilized to remove adapter sequences and trim bases with subpar quality before analysis. Following this, the refined reads were aligned to the Homo sapiens (GRCh38) genome using HISAT v2.1.0 [23], based on the HISAT and Bowtie2 implementations. The reference genome sequence and gene annotation data were obtained from the NCBI Genome assembly and the NCBI RefSeq database, respectively.

The aligned data, in SAM file format, were sorted and indexed using SAMtools v1.9. StringTie v2.1.3b was employed for the assembly and quantification of transcripts after alignment [24, 25]. Quantification was conducted at both gene and transcript levels, with results presented as raw read count, FPKM (Fragments Per Kilobase of transcript per Million mapped reads), and TPM (Transcripts Per Million). Statistical analyses of differential gene expression were performed with edgeR exactTest, using raw counts as input [26]. Hierarchical clustering was performed on rlog transformed values for the significant genes, with parameters set to Euclidean distance for the distance metric and complete for the linkage method.

### Gene set enrichment analysis (GSEA)

GSEA of RNA-seq data was conducted to investigate potential mechanisms underlying resistance to PD-1 blockade in OC. The reference gene sets were obtained from the Molecular Signatures Database (MSigDB). This included the following categories: Hallmark (H), Curated (C2), Ontology (C5), Immunologic Signature (C7), and Cell Type Signature (C8) gene sets [27]. Significance was determined by nominal p-values less than 0.05 and a false discovery rate (FDR) less than 0.25.

### CIBERSORTx analysis

The relative proportions of 22 types of tumor-infiltrating immune cells were analyzed using the CIBERSORTx algorithm, in conjunction with the LM22 matrix [28, 29]. The normalized RNA-seq gene expression file of the tumors in Hu-mice was uploaded to the CIBERSORTx portal (https://cibersortx.stanford.edu/) as a mixture file. The LM22 gene signature was then employed to run the algorithm, using 100 permutations. The resulting relative fractions of the 22 immune cell types were visualized in a heatmap and a bar chart.

### Statistics

Statistical analysis was performed using GraphPad Prism software. Either a Student’s t-test or analysis of variance (ANOVA) was used for comparison.

## RESULTS

### Tumor growth of ovarian cancer cells in Hu-mice following intraperitoneal injection

To confirm the establishment of the intraperitoneal xenograft model of OC in Hu-mice, we intraperitoneally inoculated Hu-mice with two luciferase-expressing cell lines (the highly invasive SKOV-3 Luc and the less invasive OVCAR-3 Luc [30]) after humanization via engraftment of CD34^+^ cells. We then monitored the weekly tumor growth progression over time using bioluminescence imaging (BLI) (Figure 1A). Both models showed growth in Hu-mice, with SKOV-3 Luc demonstrating more aggressive growth than OVCAR-3 Luc (Figure 1B-E).

**Figure 1.**
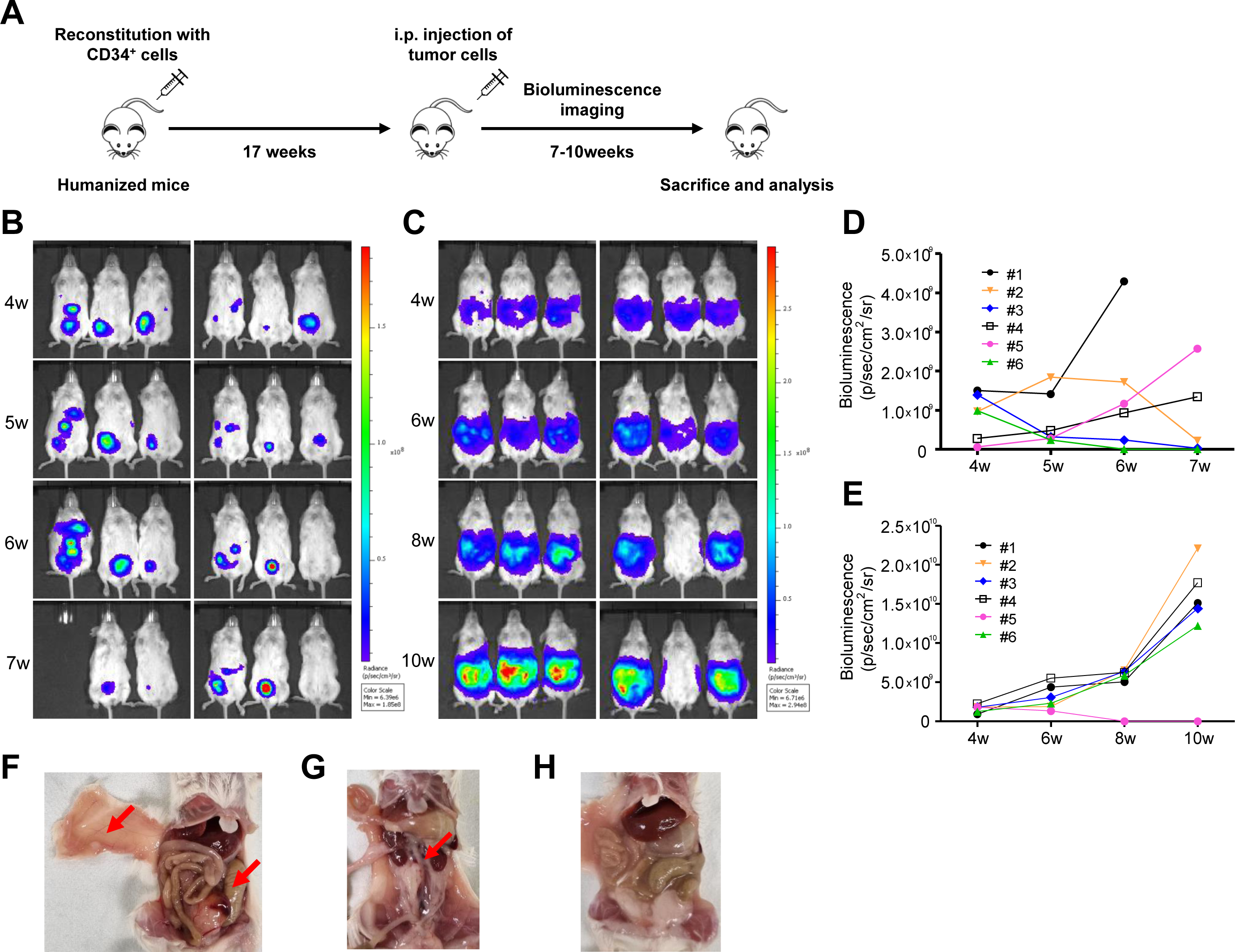
Growth of ovarian cancer cells as intraperitoneal xenografts in Hu-mice. (A) A schematic overview of the experimental setup for generating intraperitoneal ovarian cancer xenograft model in Hu-mice. Briefly, 1 × 10^7^ SKOV-3 Luc and OVCAR-3 Luc cells were injected intraperitoneally into Hu-mice, respectively, at 17 weeks after transplantation of human CD34^+^ cells. In vivo bioluminescence images (BLI) were repeatedly obtained until 7-10 weeks after IP injection of (B, D) SKOV-3 Luc and (C, E) OVCAR-3 Luc cells. Increase in BLI intensity of photons over time demonstrated growth of tumors. After 7-10 weeks post-transplant, (F) representative images of tumors (red arrow) from Hu-mice injected with SKOV-3 Luc and (G) OVCAR-3 Luc cells. (H) Representative image of milky ascites formation in OVCAR-3 Luc model.

All mice in the SKOV-3 Luc model with increasing BLI exhibited solid tumor tissues in the intraperitoneal cavity. In contrast, the mice in the OVCAR-3 Luc model developed milky ascites (Figure 1F-H). The tumor take rates in the SKOV-3 Luc and OVCAR-3 Luc models were 50.0% and 8.3%, respectively, and the rates of ascitic fluid formation were 25.0% and 83.3% within 7-10 weeks, respectively.

### Immunological characterization of ovarian xenograft tumors in Hu-mice

Tumor-infiltrating immune cells, such as lymphocytes and myeloid cells, along with the expression of programmed cell death ligand 1 (PD-L1), have been associated with the response to PD-1/PD-L1 ICI treatment [31]. We performed colorimetric immunostaining to identify the presence of TILs and PD-L1 expression in the tumors of Hu-mice. We observed CD3^+^, CD4^+^, and CD8^+^ human T cells within both SKOV-3 Luc and OVCAR-3 Luc xenografts derived from these Hu-mice (Figure 2A and B). Additionally, we demonstrated distinct PD-L1 expression on the tumors (Figure 2B).

**Figure 2.**
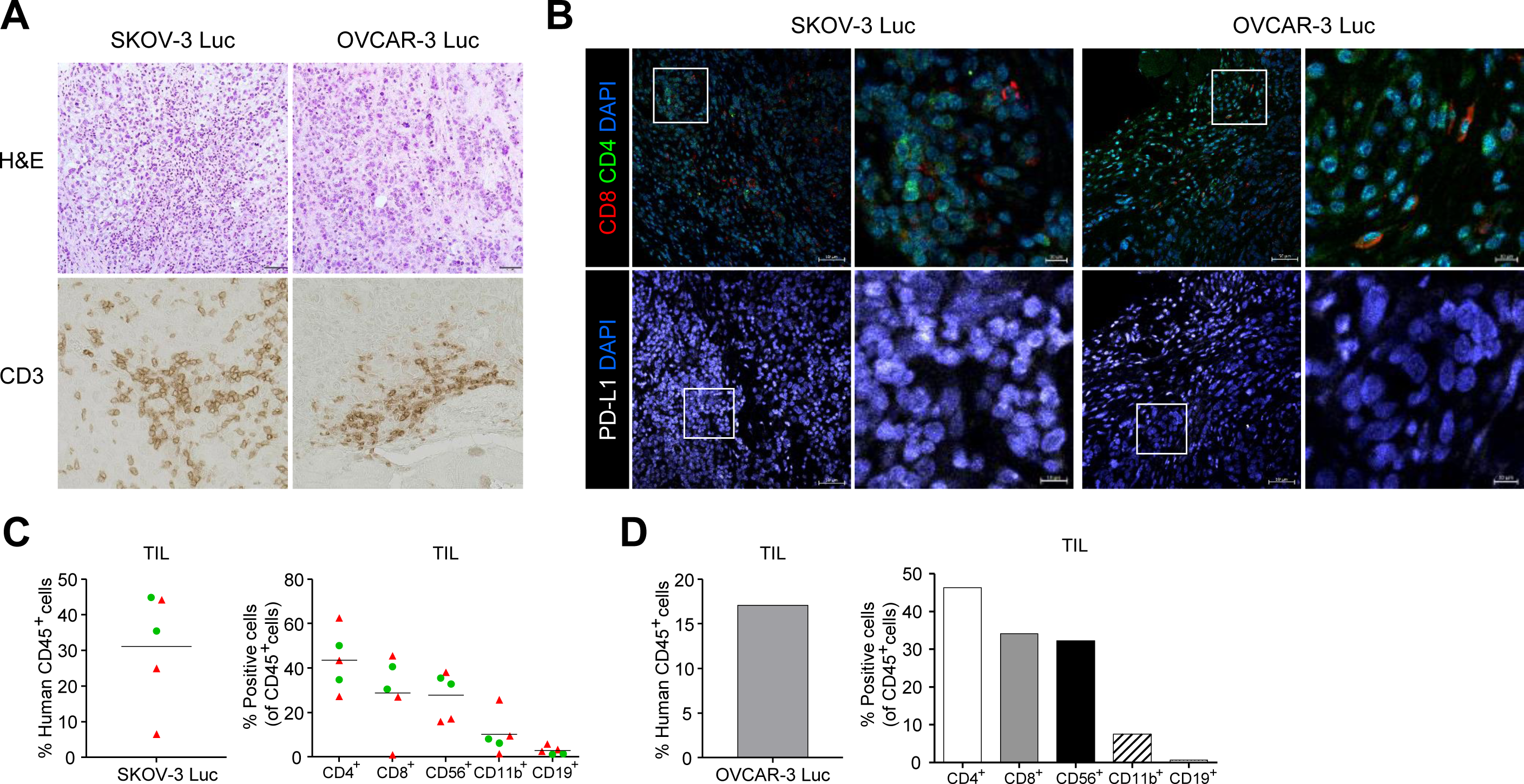
Presence of tumor-infiltrating lymphocytes (TILs) and immunophenotypical characterization of tumors in Hu-mice injected with ovarian cancer cells. The infiltration of human lymphocytes and the immunophenotype within the ovarian xenograft tumors of Hu-mice were determined by colorimetric immunostaining and flow cytometry analysis. Tumor samples were collected from the peritoneal cavity, processed through formalin fixation and paraffin embedding, then sectioned for histologic examination and analysis of immune cells and PD-L1 expression. (A) The tumor characteristics are visualized via hematoxylin and eosin staining (H&E, top panels) and immunohistochemical staining of CD3^+^ T cells (bottom panels) in tumors. (B) Immunofluorescence imaging enabled the detection of CD4^+^ and CD8^+^ T cells by utilizing anti-human CD4 (Green) and CD8 (Red) antibodies (top panels). PD-L1 expression was determined using the anti-human PD-L1 (White) antibody (bottom panels). DAPI was used for nuclear staining (Blue). Magnification, 20×; Scale bar, 50 μm. Solid tumor tissues from Hu-mice injected with ovarian cancer cells were dissociated into single cells using enzymes and mechanical methods. These collected cells were then labeled with fluorochrome-conjugated anti-human antibodies. The presence and frequency of human CD45^+^ cells, CD4^+^ T cells, CD8^+^ T cells, CD56^+^ natural killer (NK) cells, CD11b^+^ myeloid cells, and CD19^+^ B cells within the tumor tissues of (C) SKOV-3 Luc and (D) OVCAR-3 Luc models were evaluated using flow cytometry. The first batch was denoted by red triangles, and the second batch was denoted by green circles.

Concurrently, flow cytometry analysis revealed the presence of not only human lymphocytes but also myeloid cells within the tumors of SKOV-3 Luc and OVCAR-3 Luc in Hu-mice. More specifically, we observed human CD45^+^ hematopoietic cells, CD4^+^ T cells, CD8^+^ T cells, CD56^+^ NK cells, CD11b^+^ myeloid cells, and CD19^+^ B cells.

The SKOV-3 Luc models presented a range of tumor-infiltrating human CD45^+^ cells, ranging from 6.5% to 44.9%, with an average of 32.7%. Within this CD45^+^ cell population, the frequency of CD4^+^ and CD8^+^ T cells spanned 27.3% to 62.6% and 0.9% to 45.6%, averaging at 43.4% and 30.1%, respectively. Moreover, the prevalence of CD56^+^ cells within CD45^+^ cells varied from 15.9% to 38.1% (mean = 29.0%), and CD11b^+^ cells ranged from 1.4% to 25.7% (mean = 9.7%). In contrast, the average frequency of CD19^+^ B cells was quite low at 2.6%, with a range from 1.2% to 5.9% (Figure 2C).

In the OVCAR-3 Luc xenograft tumor, the proportion of tumor-infiltrating human CD45^+^ cells was 17.1% of the viable cells within the tumor. Within the CD45^+^ cell subset, the percentages of CD4^+^ and CD8^+^ T cells varied, comprising 46.3% and 34.2% of human CD45^+^ cells, respectively. Additionally, the frequencies of CD56^+^ cells and CD11b^+^ cells were observed at 32.3% and 7.6%, respectively. As with the SKOV-3 Luc model, CD19^+^ B cells in the OVCAR-3 Luc tumor remained extremely low, accounting for up to only 0.7% (Figure 2D).

### Immunophenotyping of draining lymph node, peripheral blood, and spleen in ovarian tumor-bearing Hu-mice

We next conducted an immune profiling analysis of tumor-draining lymph nodes (DLNs), peripheral blood samples, and spleens from Hu-mice injected with SKOV-3 Luc and OVCAR-3 Luc cells. Lymph nodes in NSG mice are notably underdeveloped, as evidenced by their remarkably small size, which in some cases renders them nearly undetectable [32]. However, we successfully obtained a lymph node of sufficient size located near the SKOV-3 Luc and OVCAR-3 Luc tumor tissues in Hu-mice.

A DLN from the SKOV-3 Luc model revealed a significant proportion of human CD45^+^cells, demonstrating an immune cell subset pattern similar to that of the tumors (Figure 3A and B). In the blood and spleen, more than 40% of the human CD45^+^ cell populations were detected (Figure 3C). The proportion of CD19^+^ B cells within the CD45^+^ cell population ranged from 1.0 to 27.4% (mean = 13.1%) and from 2.9 to 51.5% (mean = 25.7%) in the blood and spleen, respectively. These values were notably higher than those observed in tumors and DLN, in contrast to the frequencies of CD4^+^, CD8^+^, CD56^+^, and CD11b^+^ cell subsets, which exhibited a similar pattern (Figure 3D-H).

**Figure 3.**
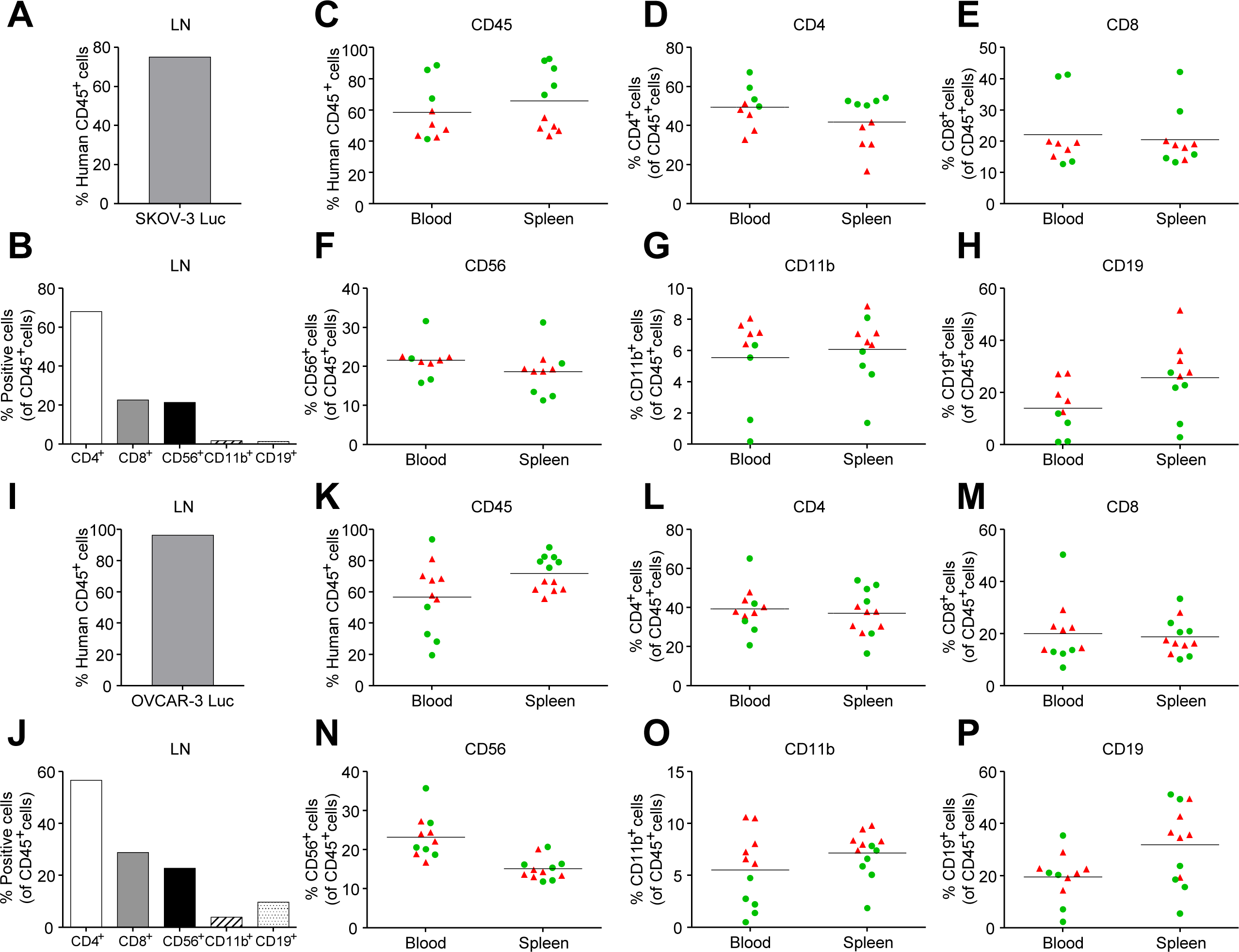
Immunophenotypical characterization of draining lymph node, blood and spleen in Hu-mice injected with ovarian cancer cells. Immunophenotyping for the prevalence of human immune cells within the draining lymph node, blood, and spleen of Hu-mice injected with (A-H) SKOV-3 Luc and (I-P) OVCAR-3 Luc cells. Flow cytometry analysis of human CD45^+^ cells, CD4^+^ T cells, CD8^+^ T cells, CD56^+^ natural killer (NK) cells, CD11b^+^ myeloid cells, and CD19^+^ B cells are shown. The first batch was denoted by red triangles, and the second batch was denoted by green circles.

The DLN of the OVCAR-3 Luc model also displayed a very high proportion of human CD45^+^ cells, reaching up to 96.3% (Figure 3I). The immune cell phenotypes of the DLN, blood, and spleen were consistent with those of the SKOV-3 Luc model, showing relatively higher frequencies of B cells (Figure 3J-P).

### Refractory response to pembrolizumab and characterization of human lymphocytes in SKOV-3 Luc tumor-bearing Hu-mice

Recently, numerous clinical trials have been conducted to incorporate ICIs targeting PD-1/PD-L1 into the treatment regimen for OC [33]. To investigate the response and underlying mechanisms related to blocking the interaction between PD-1 and PD-L1 in OC, we administered pembrolizumab, an anti-PD-1 antibody, to the intraperitoneal SKOV-3 xenograft model in CD34^+^ Hu-mice. The development of visible SKOV-3 Luc tumors was monitored for two weeks using BLI. Subsequently, the tumor-bearing Hu-mice were randomly divided into two groups: PBS control and pembrolizumab. These mice were then treated for three weeks either with PBS or pembrolizumab, depending on their assigned group. We observed that the body weight of the mice remained relatively stable throughout the treatment (Figure S1). A comparison of tumor growth revealed no significant difference between the control and pembrolizumab groups during the treatment period (Figure 4A-D). We then determined the histopathological attributes of the tumor tissues, which were excised from the peritoneal cavity of the Hu-mice (Figure S2A and B). The volume of these tumors was calculated by selectively choosing and measuring only the distinct tumor tissues. Consistently, no substantial differences in tumor volumes were found between the two groups, suggesting that the intraperitoneal SKOV-3 Luc xenograft Hu-mouse model exhibited a refractory response to pembrolizumab treatment (Figure 4E and F).

**Figure 4.**
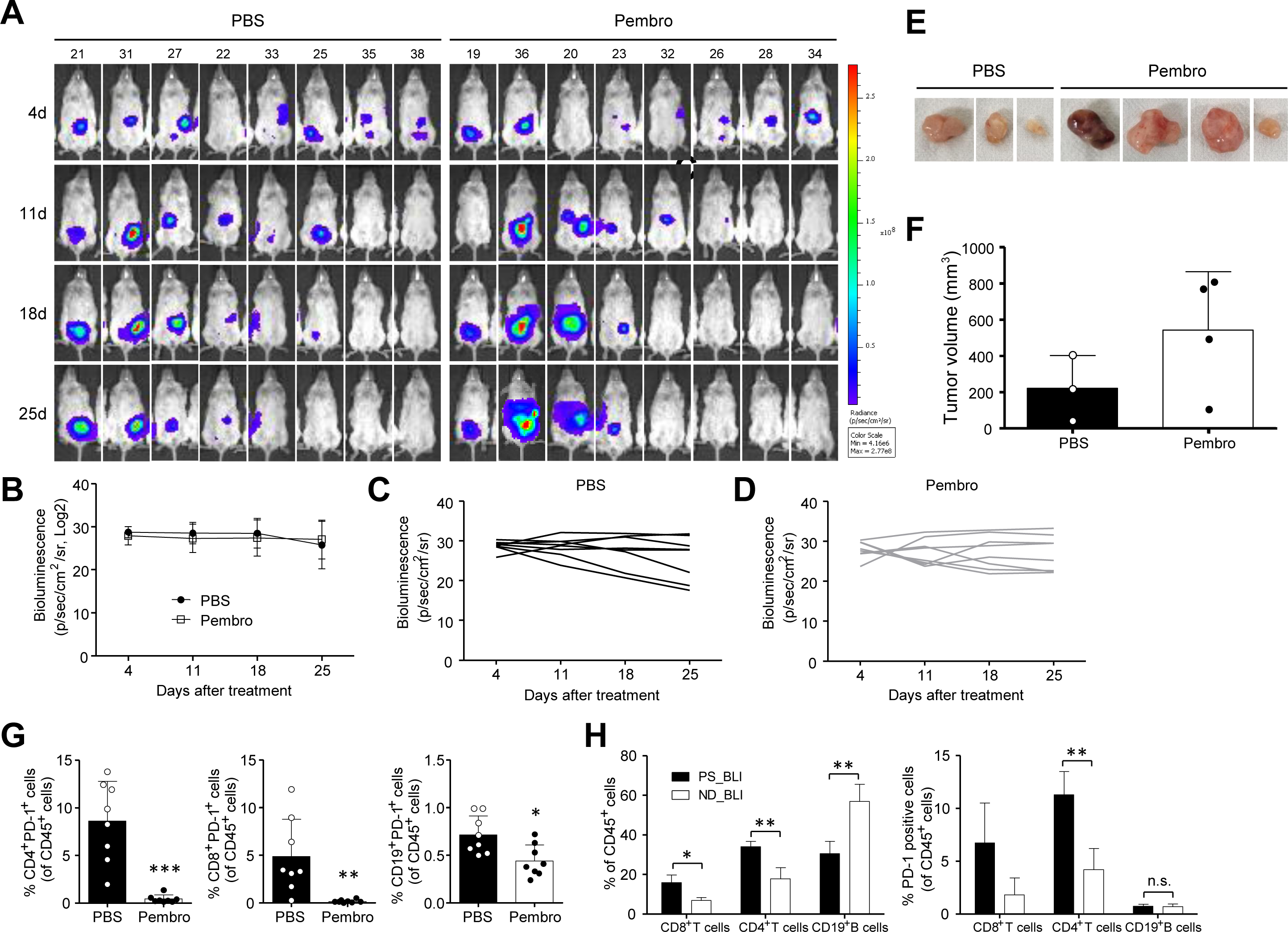
Refractory response to anti-PD-1 treatment and characterization of human lymphocytes in SKOV-3 Luc tumor-bearing Hu-mice. The sensitivity of the Hu-mice ovarian xenograft model to PD-1 blockade was investigated using SKOV-3 Luc cells, which were inoculated intraperitoneally in Hu-mice. The treatment commenced two weeks post-inoculation of cancer cells, with either a PBS control or a twice-weekly intraperitoneal dose of 10 mg/kg pembrolizumab for three weeks. (A) The BLI signal was measured at four distinct time points: 4 days, 11 days, 18 days, and 25 days post-treatment administration. (B) The intensity of BLI was quantified in (C) PBS control and (D) pembrolizumab groups. (E-F) SKOV-3 Luc cell line-derived tumors were dissected from the peritoneal cavity of Hu-mice and their volumes were subsequently measured. The effectiveness of pembrolizumab was confirmed through the analysis of target PD-1^+^ lymphocytes in Hu-mice SKOV-3 Luc xenograft model. (G-H) Frequencies of human CD4^+^ T cells, CD8^+^ T cells, and CD19^+^ B cells expressing PD-1 in the blood were evaluated using flow cytometry. Data from mice in the PBS group were then analyzed and compared between two sets: those categorized under ND_BLI and those under PS_BLI. The ND_BLI refers to instances where no BLI was detected at the end of the experiment. In contrast, PS_BLI represents instances where there was either a steady or progressively increasing exposure to BLI at the end of the experiment. P values were calculated using unpaired t-test (* P < 0.05, ** P <0.01, *** P < 0.001).

To assess the activity of pembrolizumab, a drug that binds to and saturates PD-1, we used flow cytometry to quantify human PD-1^+^ lymphocytes, including CD4^+^ T cells, CD8^+^ T cells, and CD19^+^ B cells. Previous studies have shown that depletion of human PD-1^+^ cells can be achieved using pembrolizumab or nivolumab in Hu-mice xenograft models [34, 35]. We found a significant reduction in the abundance of PD-1^+^ lymphocytes in the peripheral blood of Hu-mice treated with pembrolizumab compared to the PBS control group (Figure 4G). Analysis of the frequency of PD-1^+^ lymphocytes revealed a higher abundance of both CD8^+^ and CD4^+^ T cells in SKOV-3 Luc tumor-bearing Hu-mice compared to mice without detectable BLI and tumor growth. In contrast, the frequency of CD19^+^ B cells was lower in tumor-bearing mice. The increased prevalence of these T cells is consistent with a higher frequency of PD-1^+^ T cells. However, no observable difference in the frequency of PD-1^+^ B cells was found (Figure 4H).

### Composition of immune cells in tumor tissues of pembrolizumab-unresponsive Hu-mice

To investigate the changes in the TME of unresponsive SKOV-3 Luc-derived tumors treated with pembrolizumab in Hu-mice, we examined and compared the transcriptomes of these tumors with those of the PBS control group. We identified 2,324 differentially expressed genes (DEGs) with |log2 fold change (FC)| > 2 and p < 0.05, as illustrated in the volcano plot in Figure 5A. Among these DEGs, 814 were upregulated and 1,510 downregulated in tumors of Hu-mice treated with pembrolizumab.

**Figure 5.**
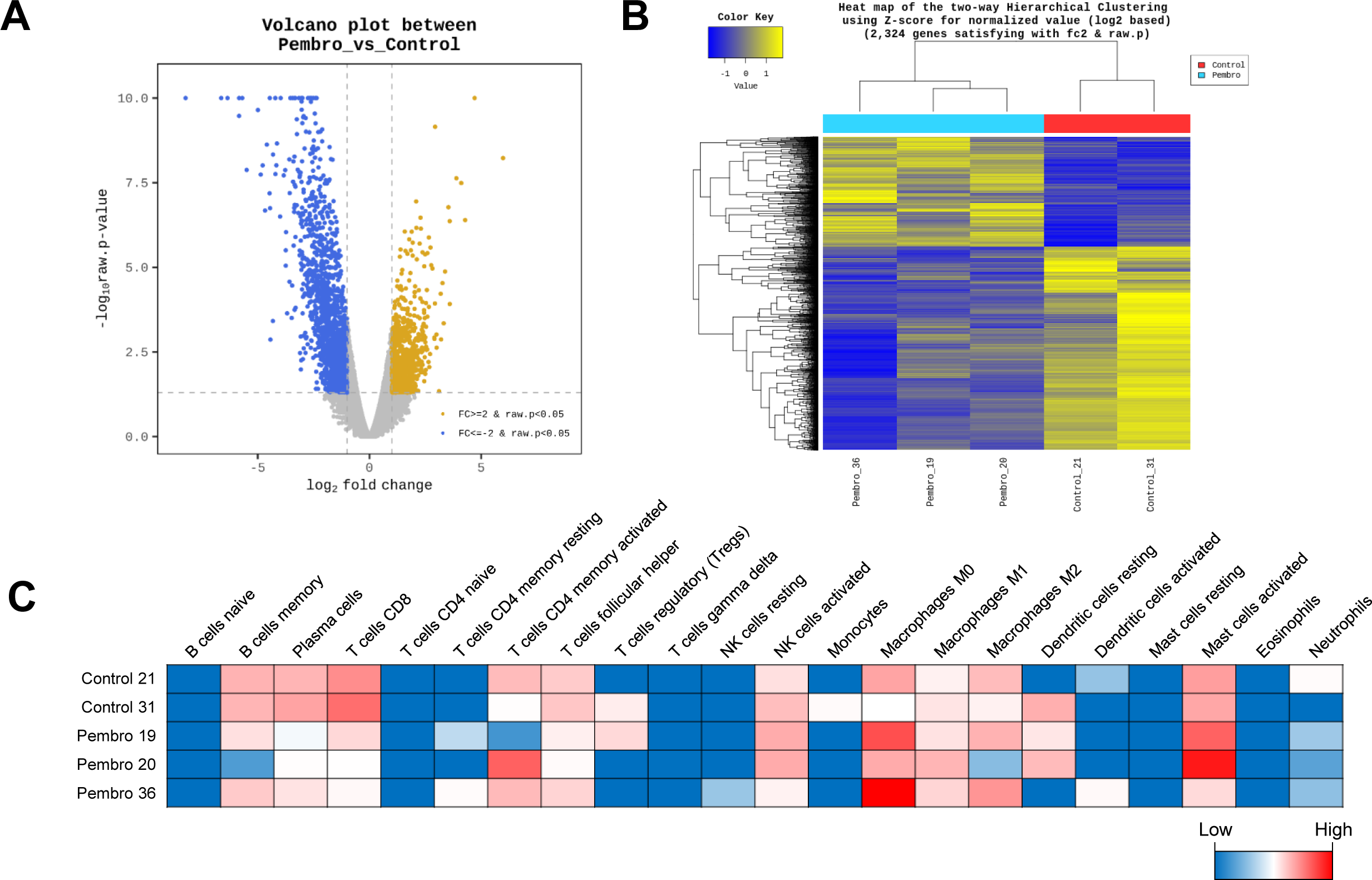
Transcriptome profiles and CIBERSORTx analysis of RNA sequencing data in tumor tissues of pembrolizumab non-responsive and PBS-controlled Hu-mice. The RNA-seq analysis examined the differences in gene expression within the TME of SKOV-3 Luc tumors unresponsive to pembrolizumab treatment, compared to the PBS control group. (A-B) Volcano plots and heatmap showing differentially expressed genes (DEGs, 814 upregulated and 1,510 downregulated genes). (A) Y-axis of volcano plots displays the p-value (-log 10) of the mean expression, and the x-axis shows the logL2-fold change value. Gold (upregulated genes, log2FC > 2) and Blue (downregulated genes, log2FC < -2) dots indicate significant DEGs (p < 0.05). (B) The genes (rows) and samples (columns) are clustered using Euclidean distance and complete-linkage methods. The heatmap displays z-scores for log2-normalized read counts that are statistically significant (|log2(FC)| > 2, p < 0.05). Yellow color indicates higher gene expression and blue color indicates lower gene expression. (C) The 22 subtypes of tumor-infiltrating immune cells are quantified using CIBERSORTx analysis. The heatmap visually represents the distribution of immune cell compositions in SKOV-3 Luc xenograft tumors, comparing those treated with PBS and Pembrolizumab.

To further assess the similarity of gene expression patterns between the tumor samples, we employed heatmap-based hierarchical clustering using Euclidean distance and complete-linkage methods (Figure 5B). Additionally, we utilized CIBERSORTx analysis along with the LM22 signature matrix to examine the relative proportions of different immune cell subpopulations in the RNA-seq data. This digital cytometry method elucidates the proportions of 22 distinct tumor-infiltrating immune cell subtypes in SKOV-3 Luc tumors of Hu-mice that were nonresponsive to pembrolizumab and subsequently compared these findings with PBS control mice (Figure 5C and S3).

Upon analysis, macrophage fractions were found to be highly abundant within SKOV-3 Luc tumors from Hu-mice, consistent with previously reported CIBERSORT results from the RNA-seq data of The Cancer Genome Atlas (TCGA) for OC [36]. Moreover, pembrolizumab-refractory xenograft tumors exhibited reduced fractions of CD8^+^ T-cells, memory B cells, and plasma cells, as well as increased proportions of naïve M0 macrophages and activated mast cells, compared to PBS control tumor tissues.

### Identifying gene signatures associated with anti-PD-1 resistance in SKOV-3 Luc xenografts of Hu-mice

We performed a gene set enrichment analysis (GSEA) on the RNA-seq data from SKOV-3 Luc xenografts in Hu-mice to examine the molecular profile of cancer immunity associated with resistance to anti-PD-1 therapy. Remarkably, we found 39 significantly enriched gene sets in pembrolizumab-nonresponsive tumors that were upregulated compared to the PBS control group (Figure 6A). The differentially expressed genes (DEGs) within these sets were predominantly enriched in RB1 target genes, KLF1 and 3 target genes, genes associated with epithelial-mesenchymal transition (EMT) and fibroblasts, as well as TCF21, MYD88, and SMARCE1 target genes (Figure S4). Furthermore, we identified 4 significantly enriched gene sets among the downregulated DEGs, encompassing interferon-gamma and alpha response (Figure 6B).

**Figure 6.**
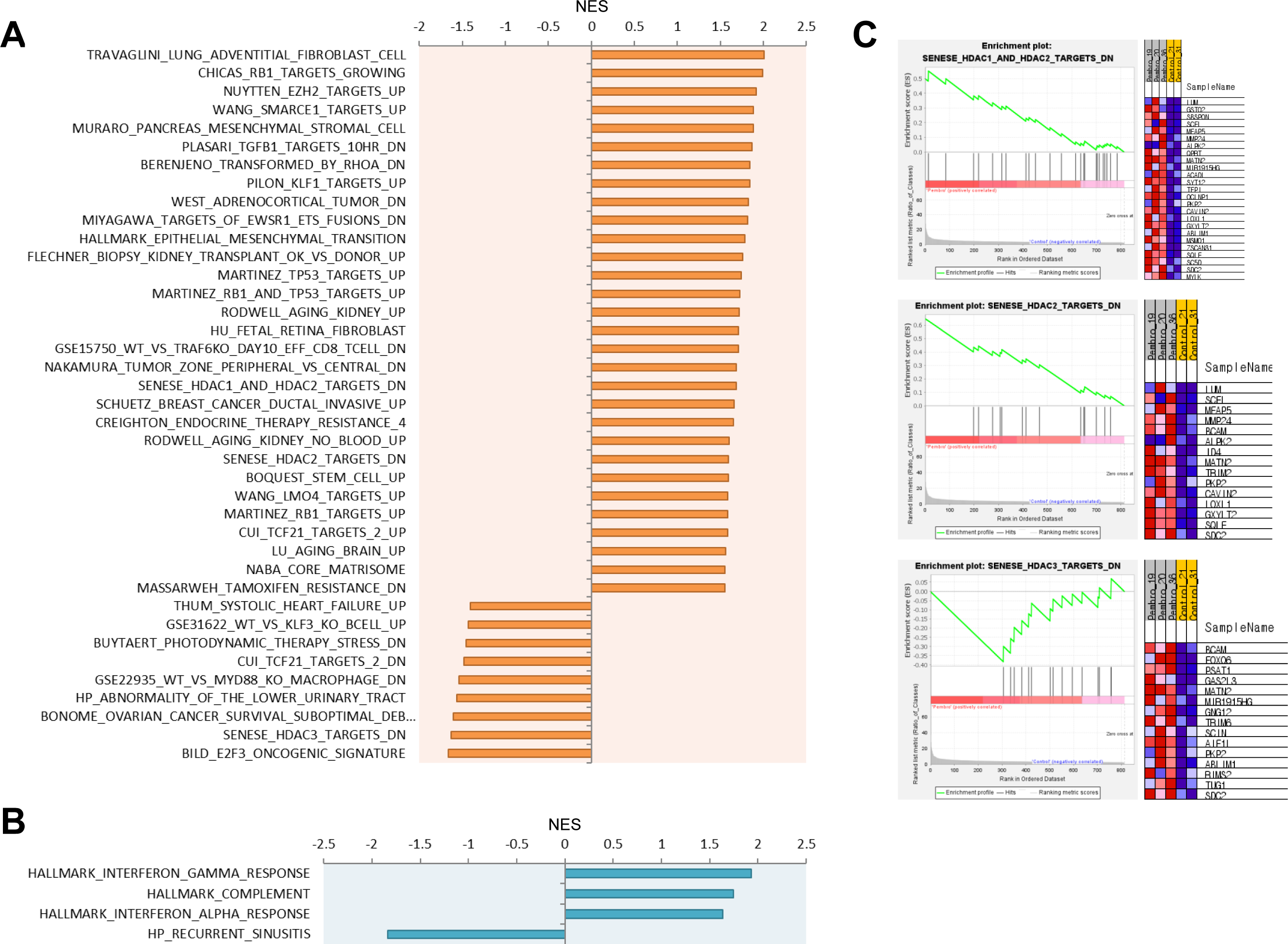
Gene set enrichment analysis (GSEA) results in pembrolizumab refractory xenograft tumors. The gene signatures in pembrolizumab non-responsive SKOV-3 Luc tumors were identified by GSEA. Significantly enriched gene sets (FDR q-value < 0.25) among (A) upregulated and (B) downregulated genes in the pembrolizumab compared to the control group. (C) Enrichment plots show the genes associated with histone deacetylase (HDAC) targets. All gene sets were selected from the molecular signatures database (MSigDB), specifically categories C2, C5, C7, C8, or H.

Notably, one of the main gene sets upregulated in pembrolizumab-nonresponsive tumors was found to be the histone deacetylase (HDAC) class I target genes (Figure 6A and C). Previous studies have revealed the crucial role of HDAC in cancer immunotherapy. HDAC inhibitors are currently under active clinical examination and are being combined with immunotherapy for solid tumors, a strategy aimed at amplifying therapeutic responses [37].

## DISCUSSION

Preclinical studies using animal models that accurately mimic the human immune system are crucial in elucidating the potential clinical outcomes and underlying mechanisms of immuno-oncology (IO) drugs [38, 39]. These studies aid in understanding the TME and assessing the prognostic significance of IO drugs, such as PD-1/PD-L1 ICIs, which have shown limited efficacy in treating OC [17, 35, 40]. In this study, we demonstrated the tumor growth of two intraperitoneal OC xenografts in Hu-mouse models and characterized their morphological and immunophenotypic features. Furthermore, we identified several pivotal genes and biological pathways in the TME of OC that could serve as potential therapeutic targets for overcoming anti-PD-1 treatment resistance. This was achieved using SKOV-3 Luc humanized xenografts that showed non-responsiveness to pembrolizumab.

In recent years, significant progress has been made in the development and application of Hu-mouse xenograft models. The most common methods for testing cancer immunotherapy involve the subcutaneous implantation of established cell lines and patient-derived tissues [16, 18, 35, 41]. However, these models may not always accurately reflect the characteristics of the primary tumor, as they lack interaction with the TME, and do not provide the conditions necessary for metastasis. Orthotopic xenografts can offer a more accurate representation of the metastatic behavior of tumors and their TME [42, 43]. Previous studies have explored the development of orthotopic xenograft models in Hu-mice, using luciferase-positive human pancreatic and osteosarcoma cells [44, 45]. The orthotopic OC xenograft model in Hu-mice, using the OV-90 luciferase cell line and PDX cells, was initially employed to verify the response to nivolumab [19]. As an alternative approach that avoids surgical incisions and offers simplicity, we intraperitoneally injected the luciferase-expressing SKOV-3 and OVCAR-3 cell lines into CD34^+^ Hu-mice. We confirmed tumor formation and progression through in vivo BLI (Figure 1B-D). In Hu-mice, both SKOV-3 Luc and OVCAR-3 Luc intraperitoneal xenograft models were less aggressive and showed a relatively long survival time (more than 6 weeks), consistent with previous studies [20, 46].

Tumor-infiltrating immune cells play a crucial role in modulating responses to immunotherapy in many solid tumors. The response to anti-PD-1/PD-L1 therapy is positively correlated with the degree of pre-existing TILs, including CD8^+^ T cells expressing PD-L1 [7]. Conversely, immunosuppressive immune cells such as tumor-associated macrophages (TAMs) and regulatory T cells (Tregs) have been associated with a poor prognosis in ovarian cancer (OC) [47-49]. These cells facilitate tumor immune evasion, potentially leading to resistance to PD-1/PD-L1 therapy [50]. In our study, we characterized the profiles of human immune cells within tumors, DLNs, blood, and spleen of Hu-mice. Our observations confirmed the presence of human immune subsets involved in both lymphocytes and myeloid cells, which are responsible for innate and adaptive immune responses. We identified a significant proportion of CD11b^+^ myeloid cells in ovarian xenografts from the peritoneal cavity of Hu-mice. These findings are consistent with other models using Hu-mice, including subcutaneous xenografts for various types of cancer and the orthotopic OC model [16, 19]. Furthermore, our transcriptome data within SKOV-3 Luc tumors of Hu-mice revealed various populations of human immune cells, including T and B lymphocytes, natural killer (NK) cells, M0/M1/M2 macrophages, dendritic cells, neutrophils, and mast cells. Interestingly, this intraperitoneal model displayed a substantial distribution of macrophages within the tumors, in contrast to subcutaneous xenografts where macrophages were found in very low proportions [16]. These results are in line with the known characteristics of the TME for patients with OC [36, 48]. Consequently, our model can be utilized to evaluate the anti-tumor efficacy of potential immuno-oncology drug treatments for OC, particularly in the TME where there is a high abundance of macrophages.

Previous studies have reported varying sensitivities in Hu-mice xenograft models exposed to PD-1/PD-L1 ICIs, such as pembrolizumab, nivolumab, and atezolizumab, across different types of cancer [16, 19, 34]. In this study, despite the effective blocking of the PD-1 protein on T and B lymphocytes when treated with pembrolizumab (Figure 4G), the SKOV-3 Luc Hu-mouse model did not show any significant inhibition of tumor growth (Figure 4A-F). Indeed, in a clinical trial, the objective response rate (ORR) of pembrolizumab as a single-agent treatment was only 8% in patients with advanced OC [10]. We observed a reduced presence of CD8^+^ T cells, memory B cells, and plasma cells in SKOV-3 Luc xenografts from pembrolizumab nonresponsive Hu-mice, compared to PBS control mice (Figure 5C). It has been reported that patients with melanoma and renal cell carcinoma who did not respond to ICIs had a lower frequency of memory B cells and plasma cells in their tumors compared to responders. These B cell subsets were associated with a more favorable response to ICIs, potentially facilitating an enhanced T cell response [51]. Interestingly, elevated levels of mast cells were observed in SKOV-3 Luc tumors that did not respond to anti-PD-1 treatment (Figure 5C). Previous research examining the anti-PD-1 response in a melanoma xenograft model using Hu-mice also reported an increase in the prevalence of tumor-infiltrating mast cells following pembrolizumab treatment. This increase is associated with resistance to PD-1 blocking, facilitated by interactions with Treg cells and the subsequent down-modulation of MHC class I [17].

The intraperitoneal CDX Hu-mouse model in our study enabled us to explore factors associated with resistance to IO drug treatments. Our RNA-seq results identified gene sets linked to anti-PD-1 resistance in OC tumors in Hu-mice unresponsive to pembrolizumab (Figure 6A and B), with a notable enrichment of HDAC class I target genes. (Figure 6C). HDAC inhibition enhances T-cell mediated anti-tumor immune responses by increasing the presence of tumor-infiltrating CD8^+^ T cells and molecules involved in antigen processing and presentation, while also blocking Treg and myeloid-derived suppressor cells (MDSCs) infiltration [52]. Additionally, our subsequent GSEA of upregulated DEGs revealed a connection between SKOV-3 Luc tumors refractory to anti-PD-1 therapy and EMT, fibroblast-related signatures, and TCF21 target genes (Figure S4D-H). EMT and cancer-associated fibroblasts (CAFs) are known to play an immunosuppressive role in the TME, contributing to tumor progression and immunotherapy resistance in several cancer types, and a significant correlation exists between EMT-related gene expression and stromal components [53]. The expression of TCF21 influences the state of CAFs. Overexpression of TCF21 affects the protoplasmic properties of fibroblast-activated protein (FAP)-high CAFs, which in turn suppresses OC growth, invasion, and metastasis [54]. Moreover, a study has shown that the mesenchymal subtype, characterized by a high presence of EMT-related gene signatures, has a lower number of intraepithelial CD8^+^ tumor-infiltrating T cells, and is associated with poor prognosis [55]. These characteristics of the subtype were consistent with those found in SKOV-3 Luc tumors in Hu-mice that did not respond to pembrolizumab.

The CD34^+^ Hu-mouse model we employed includes most cells of the human immune system. However, functional deficiencies in T cells, B cells, and NK cells have been identified, as previously reported [56]. Additionally, while achieving HLA matching between donor HSCs and tumors poses challenges in simulating fully functional human immune systems, several studies have indicated no significant correlation between the anti-tumor responses of PD-1/PD-L1 ICIs and HLA matching in CD34^+^ Hu-mice [16, 34]. In the xenograft model, tumor take rate is crucial for experimental reproducibility, and the obtained tumor samples are required for subsequent analyses. The SKOV-3 Luc xenograft model exhibited a moderate tumor take rate, while the OVCAR-3 Luc showed over 80% tumor growth in Hu-mice but had a notably low take rate of visually recognizable tumors. It has been reported that tumor growth was not only restricted in the nasopharyngeal cancer PDX model with a humanized immune system [18], but the CD8^+^ T cells in Hu-mice also inhibited melanoma tumor growth [17]. The presence of reconstituted human immune cells may have affected tumor development in our OC models. Surgical transplantation techniques, though more intricate, yielded higher take rates and faster growth compared to intraperitoneal methods [57], suggesting potential for enhancing reproducible tumor formation in future orthotopic OC Hu-mouse models. Moreover, our transcriptome results provided insights into the key factors associated with resistance to anti-PD-1/PD-L1 in OC, but our study was limited by a small sample size and the absence of tumor samples from anti-PD-1 responsive mice. To understand the underlying mechanisms more deeply, additional investigation of functional pathways of pivotal genes is necessary. Despite these limitations, our OC CDX model using CD34^+^ Hu-mice can be used as one of the powerful tools to study the efficacy and related mechanisms of human-targeted IO drugs and can provide more biological information than clinical trials.

In conclusion, our study with intraperitoneal OC CDX model using CD34^+^ Hu-mice identified potential targets for combination therapy with PD-1 ICIs to overcome resistance to PD-1 blockade in OC. The models we generated using Hu-mice can serve as valuable tools for examining the efficacy of new immunotherapies.

## Acknowledgments

We thank the core facilities of flowcytometry and optical imaging and disease animal resource center at the ConveRgence mEDIcine research cenTer (CREDIT), Asan Medical Center for the use of their shared equipment, services and expertise.

## Ethics statements

## Patient consent for publication

Not applicable.

## Ethics approval

Not applicable.

## Consent for publication

All authors have approved the manuscript and agreed with publication.

## Availability of data and material

All data relevant to the study are included in the article or uploaded as supplementary information. Data are available from the corresponding author. RNA sequencing data generated in this study are available in the GEO database with accession code GSE245913.

## Author Contributions

S.W.K., J.-Y.L., O.-J.K., and S.-W.L. designed and performed the experiments, and generated the data for the study. S.W.K., Y.-M.K., and S.-W.L. conceived and designed the study.

S.W.K. wrote the paper. S.-W.L. performed supervisory roles in conception, direction, and completion of the study. E.K.C. acquired the funding.

---

**Figure S1.**
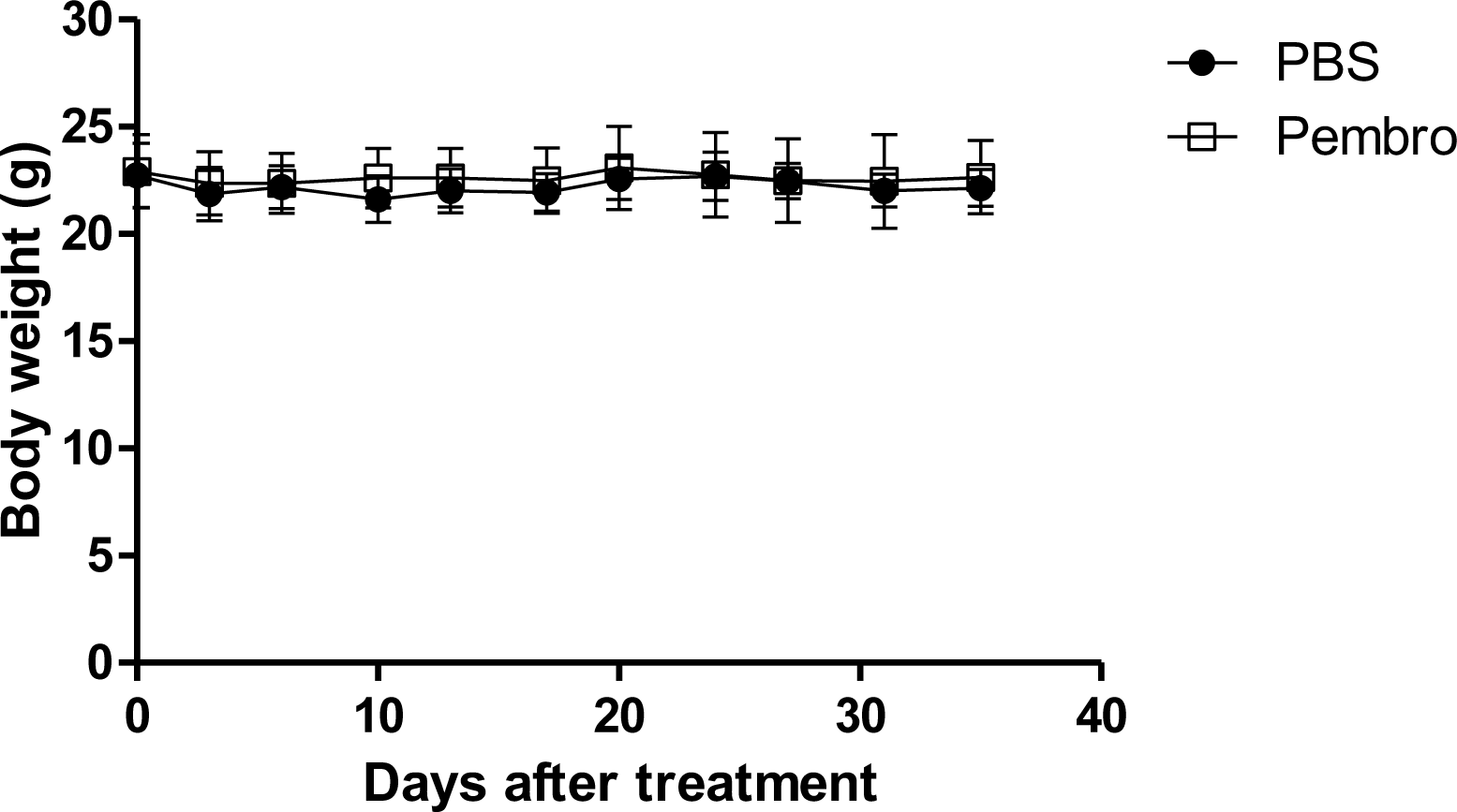
Body weight changes in Hu-mice bearing SKOV-3 Luc xenograft tumors treated with PBS or pembrolizumab. During the periods of pembrolizumab administration, the body weights of the Hu-mice remained relatively stable with no notable alterations.

**Figure S2.**
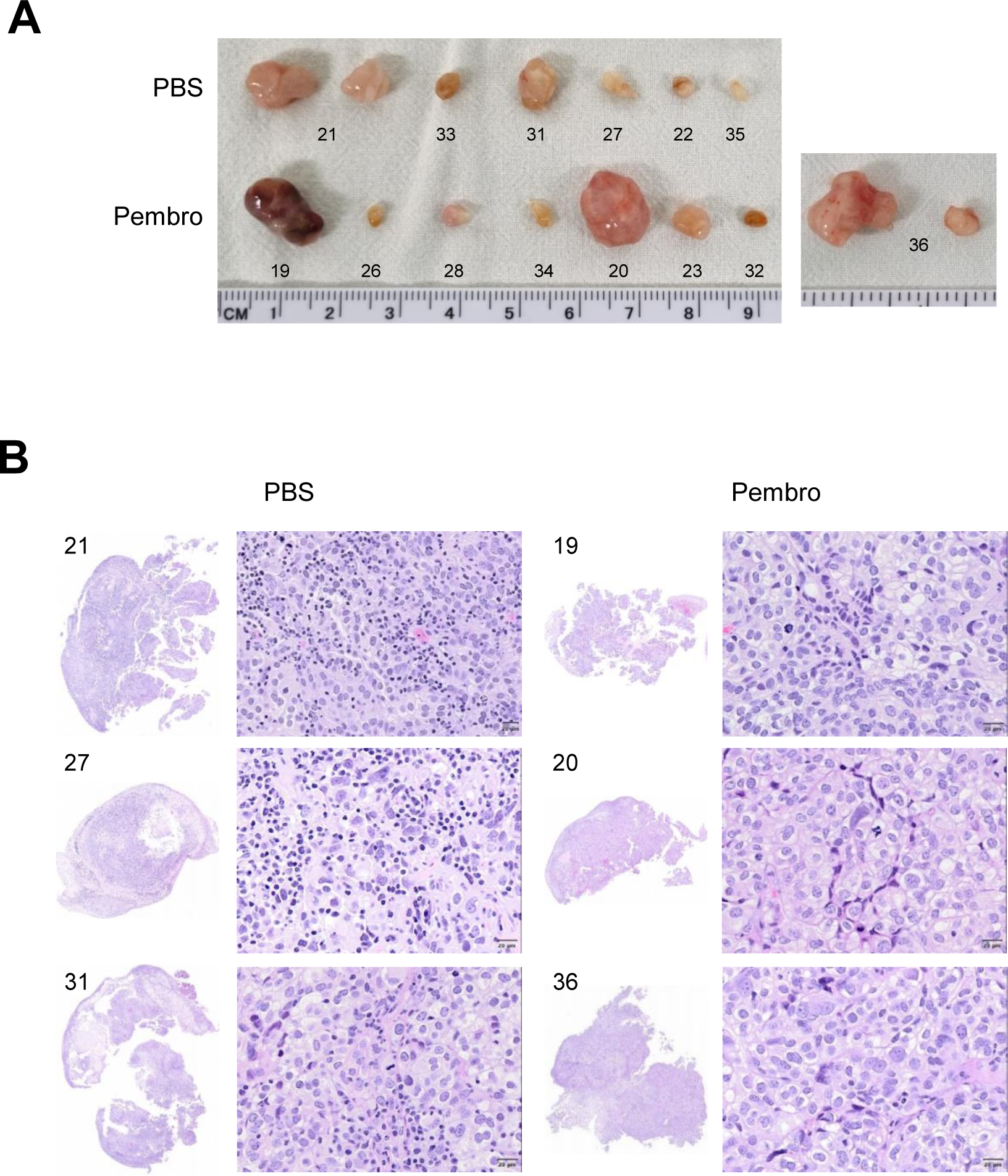
Histological examination of SKOV-3 Luc xenograft tumors in Hu-mice treated with PBS or pembrolizumab. Histopathological examination of SKOV-3 Luc xenografts in Hu-mice was performed by H&E staining. (A) Tumor tissues were resected from the intraperitoneal cavity of Hu-mice at the experimental endpoint. (B) Representative images show SKOV-3 Luc-derived tumor tissues.

**Figure S3.**
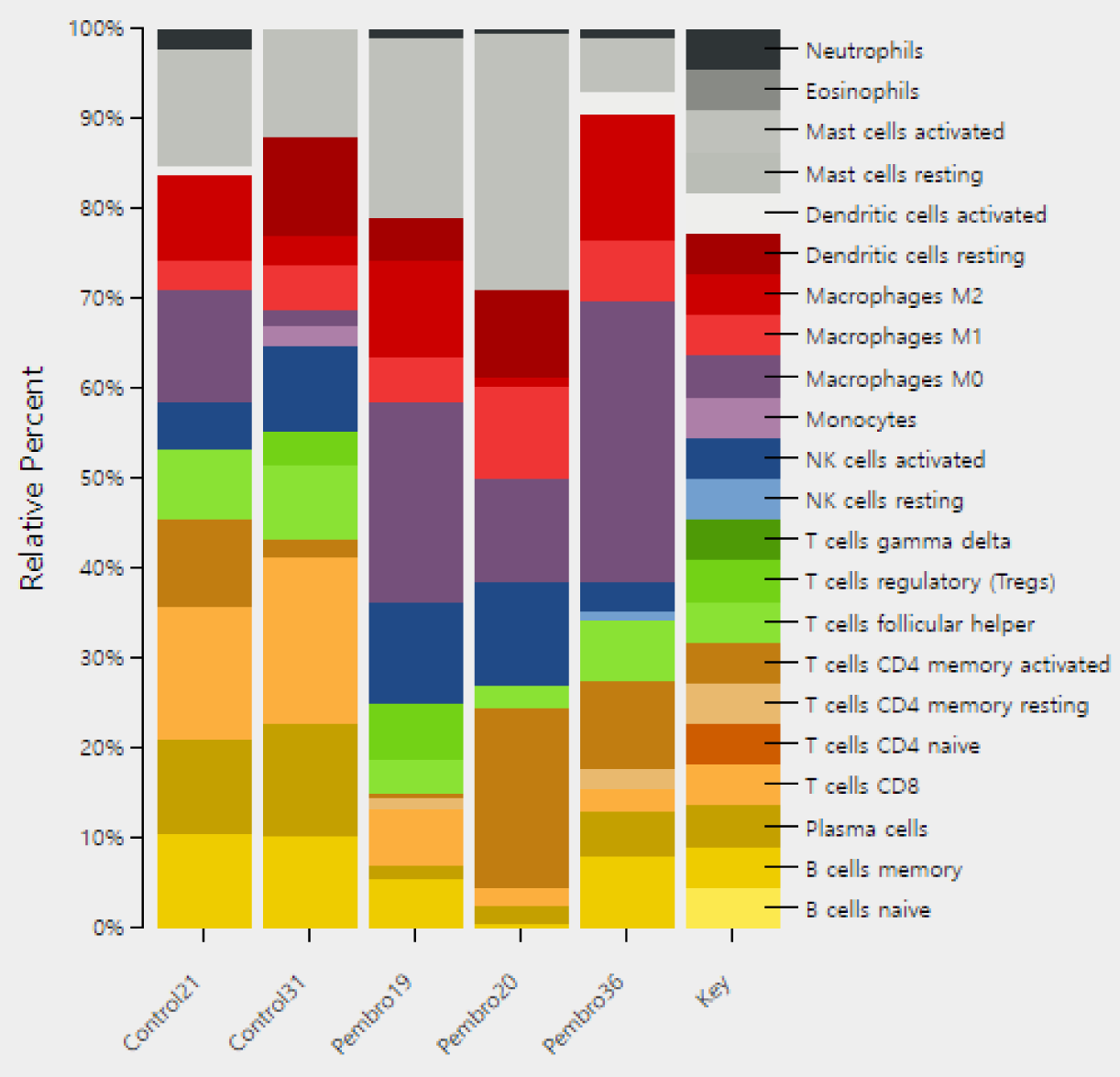
Immune cell composition in SKOV-3 Luc xenograft tumors in Hu-mice treated with PBS or pembrolizumab. The bar chart illustrates the relative abundance ratio of 22 immune cell subsets in tumors from the PBS and pembrolizumab groups. The data was obtained through CIBERSORTx using the LM22 immune cell gene signature, and the visualization was automatically created using the CIBERSORTx web server.

**Figure S4.**
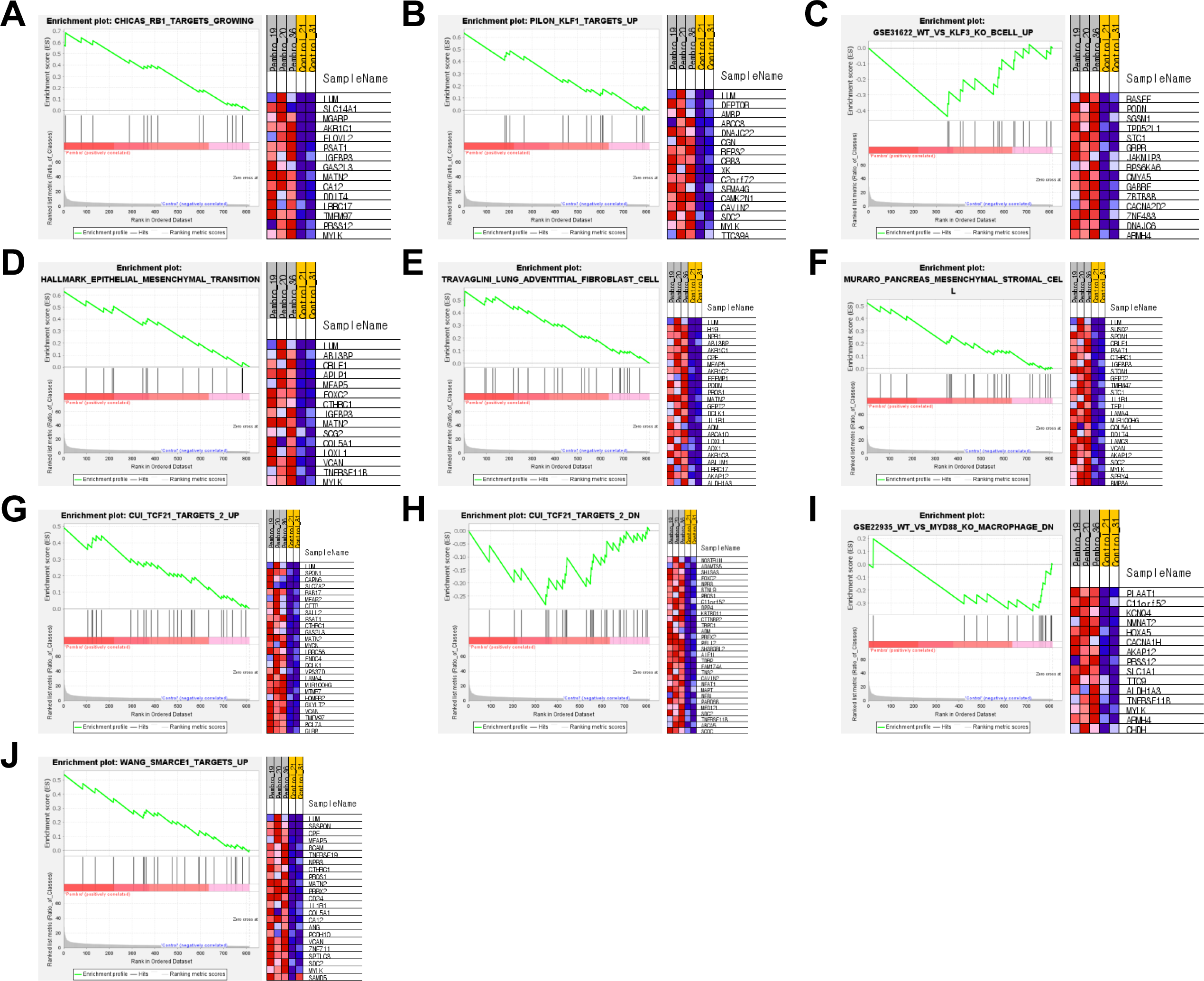
Enrichment plots from gene set enrichment analysis (GSEA) of the ten gene signatures. Representative gene signatures were significantly enriched in the SKOV-3 Luc-derived tumors of Hu-mice treated with pembrolizumab vs. PBS control (FDR < 0.25). The enrichment score (ES) is shown on the y-axis, while the x-axis represents genes (represented as vertical black lines) found in the gene sets. All gene sets were selected from the molecular signatures database (MSigDB), specifically from categories C2, C5, C7, C8, or H.

**Figure S5.**
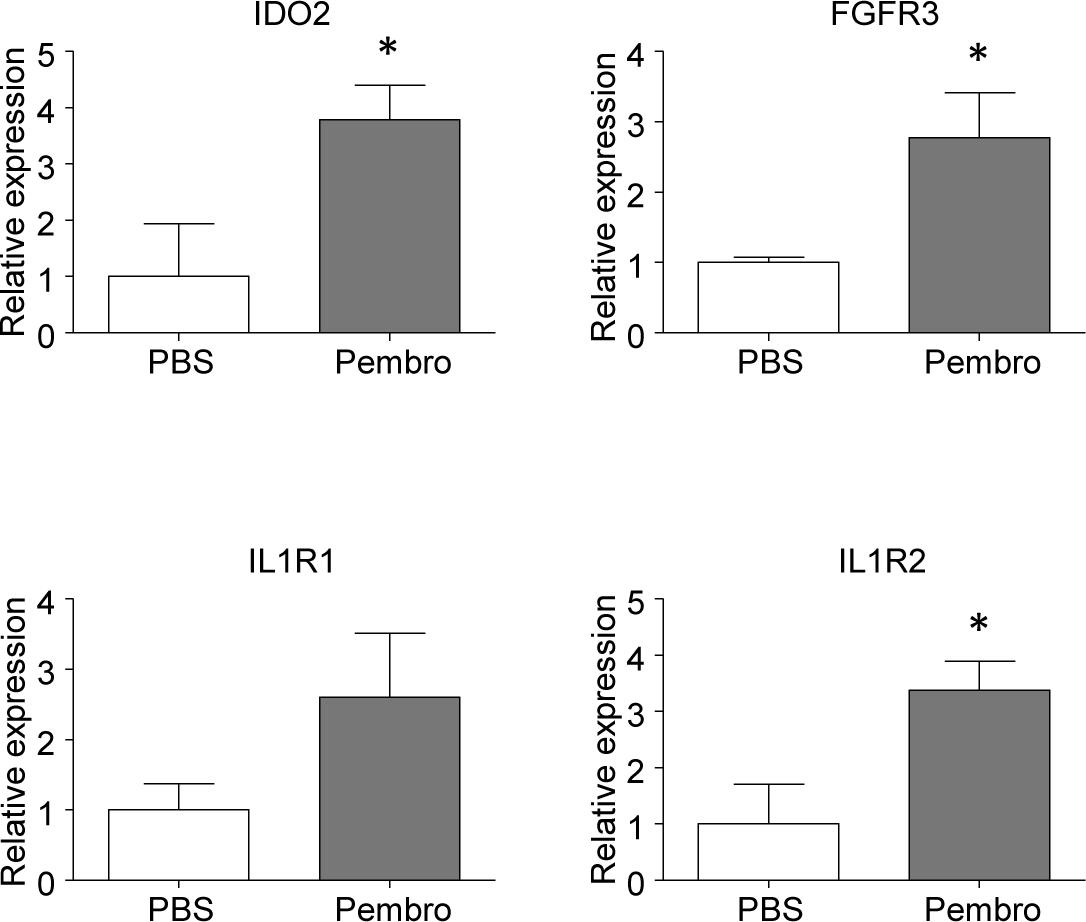
Relative expression of four genes from the RNA-seq data. The expression levels of four genes associated with cancer immunosuppression, namely IDO2, FGFR3, IL1R1, and IL1R2, are depicted. The statistical significance was determined using unpaired t-test (* P < 0.05).

